# Mechanistic home range capture–recapture models for the estimation of population density and landscape connectivity

**DOI:** 10.1101/2023.03.01.530712

**Authors:** Keita Fukasawa, Daishi Higashide

## Abstract

Spatial capture–recapture models (SCRs) provide an integrative statistical tool for analyzing animal movement and population patterns. Although incorporating home range formation with a theoretical basis of animal movement into SCRs can improve the prediction of animal space use in a heterogeneous landscape, this approach is challenging owing to the sparseness of recapture events.

In this study, we developed an advection–diffusion capture–recapture model (ADCR), which is an extension of SCRs incorporating home range formation with advection–diffusion formalism, providing a new framework to estimate population density and landscape permeability. we tested the unbiasedness of the estimator using simulated capture–recapture data generated by a step selection function. We also compared accuracy of population density estimates and home range shapes with those from SCR incorporating the least-cost path and basic SCR. In addition, ADCR was applied to real dataset of Asiatic black bear (*Ursus thibetanus*) in Japan to demonstrate the capacity of the ADCR to detect geographical barriers that constrain animal movements.

Population density and permeability of ADCR were substantially unbiased for simulated datasets. ADCR could detect envitonmental signals on connectivity more sensitively and could estimate population density, home range shapes and size variations better than the existing models. For the application to the bear dataset, ADCR could detect the effect of water body as a barrier of movement which is consistent with previous studies, whereas estimates by SCR with the least-cost path was difficult to interplet.

ADCR provides unique opportunities to elucidate both individual- and population-level ecological processes from capture–recapture data. By offering a formal link with step selection functions to estimate animal movement, it is suitable for simultaneously modeling with capture–recapture data and animal movement data. This study provides a basis for studies of the interplay between animal movement processes and population patterns.

## Introduction

Anthropogenic activity leads to habitat loss and fragmentation, posing a major threat to biodiversity. In a fragmented landscape, connectivity between habitat patches affects population patterns and sustainability (Hanski et al. 1994). Functional connectivity in a heterogeneous landscape is not only of general interest in basic ecology but also a key factor affecting decision-making in wildlife management and conservation (Goodwin and Fahrig 2002, Royle et al. 2018, Osada et al. 2019). Circuit theory and least-cost path modeling are two major paradigms for evaluating landscape connectivity, with different underlying behavioral assumptions and statistical properties (McRae et al. 2008, Marrotte and Bowman 2017). Permeability, which is a measure of connectivity based on circuit theory has explicit links with animal movement models, such as random walks and step selection functions (McRae 2006, Duchesne et al. 2015), and models of animal gene flow and movement have numerous applications (Dickson et al. 2019). Empirical determination of connectivity parameters, such as effects of land cover types on permeability, is important to improve objectivity, and the development of estimators is a major challenge in ecology (Ovaskainen 2004, Ovaskainen et al. 2008, Beyer et al. 2016, Signer et al. 2017, Osada et al. 2019).

Spatial capture–recapture models (SCRs) are powerful tools with the capacity to estimate individual movement processes and population abundance in an integrative manner (McClintock et al. 2022). An extention of SCR incorporating the least-cost path distance in the detection function instead of Euclidian distance (Royle et al. 2013, Sutherland et al. 2015, Fuller et al. 2016) can be applicable to estimate weights of landscape factors on the cost path distance. Permeability has favorable properties for practical applications, such as robustness to the resolution of landscape data (Marrotte and Bowman 2017) and a straightforward link with movement models (Diniz et al. 2020). In particular, the process of home range formation considering permeability and site fidelity has been studied extensively by step selection functions in movement ecology (Thurfjell et al. 2014, Duchesne et al. 2015, Signer et al. 2017, Péron 2019, Potts and Schlägel 2020). However, circuit theory-based SCRs are not as well-studied as SCRs incorporating the least-cost path. Although circuit theory-based SCRs can provide insight into the relationship between individual-level displacement and population-level abundance patterns, the temporal sparseness of capture–recapture data limits the application of SCR to complex movement processes (McClintock et al. 2022).

Incorporating mechanistic models of home range formation into SCR can enable inferences about permeability as well as population abundance from capture–recapture data in a computationally tractable manner. Mechanistic home range analysis (Moorcroft and Lewis 2006, Moorcroft and Barnett 2008) is an analytical approach that ties the process of animal movements to the resultant home range shape at an equilibrium state.

The approach has contributed to the development of advection–diffusion equations to obtain flexible home ranges and likelihood-based modeling to estimate parameters empirically. The advection–diffusion equations describe deterministic drift to a focal point (e.g., an activity center) as advection terms and random walks as diffusion terms. Circuit theory is analogous to a diffusion process with heterogeneous diffusivity (Hanks and Hooten 2013, Yamaura et al. 2022), and the advection–diffusion equations with heterogeneous diffusion coefficients in space are applicable to home ranges formed by animal movement affected by barriers in a landscape. Although closed-form analytical solutions of advection–diffusion equations rarely exist, computationally efficient ways to obtain an equilibrium solution have been developed (Hirsch 2007). By integrating mechanistic home range analysis into SCR, the simultaneous estimation of permeability and population density is possible, without substantial computational costs. Hereafter, we refer to this new SCR approach as the advection–diffusion capture–recapture model (ADCR).

Although ADCRs and SCRs with the least-cost path (Royle et al., 2013; Sutherland et al., 2015; Royle et al., 2018) differ with respect to connectivity assumptions and cannot be directly compared, evaluations of the performance of estimators are helpful for users. Applying statistical models to simulated data generated from a mechanistic animal movement model is a straightforward way to test the unbiasedness of estimators and robustness to deviation from assumptions (Dupont et al. 2022, Theng et al. 2022). Also, an application to real dataset is important to illustrate how statistical models can yield realistic estimates even in situations where there is unknown heterogeneity in the data generating process.

The aim of this study was to develop a formal framework for ADCR and to compare the accuracy of the estimated abundance and home range shape with those obtained by SCRs with the least-cost path. The identification of the optimal model is outside of the scope of this paper. Rather, our aim is to establish the merit of the ADCR under animal movement modeled by a step selection function, which is common in movement ecology. We also compared density estimates with those of basic SCR (Borchers and Efford 2008) as a null model of heterogeneous connectivity. We predict that all three methods, ADCR, an SCR with the least-cost path, and basic SCR, will accurately estimate population densities because SCR is robust to the violation of home range assumptions (Efford 2019). We also predict ADCR will have higher predictive ability of home range shapes than SCR with the least-cost path and basic SCR when animal movement is driven by exploratory behavior in landscapes with heterogeneous permeability. As an example of model application, we applied ADCR to estimate factors affecting peameability for Asiatic black bear (*Ursus thibetanus*) in Japan.

## Material and Methods

### Model Framework

ADCR is an extension of SCR in which animal home ranges are modeled by equilibrium solutions of the advection–diffusion equation, an Eulerian representation of stochastic animal movement in a heterogeneous connectivity landscape. By integrating the advection–diffusion equation with SCR, it is possible to simultaneously estimate population density while inferring the effect of landscape connectivity on the skewed shape of the home range, using capture–recapture data. Here, we describe the model structure of ADCR.

#### Detection model and population process

In stationary SCR modeling, a detection model that describes the encounter of an individual to a detector reflects the home range behavior of the animal. The home range of individual is defined by its utilization distribution, which is a bivariate probability distribution of an individual location in a space. In general, the model of utilization distribution has latent variables of an activity center μ = (μ*_x_*, μ*_y_*), where an individual is attracted. For example, a basic SCR (Borchers and Efford 2008, Royle et al. 2009) often assumes a half-normal detection function, which implies that the utilization distribution is a stationary bivariate normal distribution centered to the activity center. To introduce the ADCR, which covers a more flexible class of utility distributions under heterogeneous permeability landscapes, we adopted a detection model that describes an explicit link between the encounter frequency of an individual to a detector and the utilization distribution *p\**(**s** | μ), which defines the probability density at location **s** = (*x*, *y*) of an individual with an activity center μ. Note that the integration of *p\**(**s** | μ) over **s** equals 1 because of the nature of probability distribution.

An SCR survey is conducted using an array of detectors in space. Researchers may either iterate the survey on more than one occasion or treat the entire survey period as a single occasion. When the SCR data are collected by count detectors, the encounter frequency of the *i*th individual to the *j*th detector on the *k*th occasion is written as

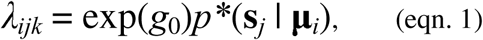

where *g*_0_ is a parameter of log detectability, μ*_i_* is the activity center of the *i*th individual, and **s***_j_* is the location of the *j*th detector. The corresponding observed value, *Y_ijk_*, follows a probability distribution depending on λ*_ijk_*, denoted by *p*(*Y_ijk_* | μ*_i_*, θ), where θ is a vector of parameters that affects λ*_ijk_*, including *g*_0_ and the parameters controlling *p\**. The number of detections follows a Poisson distribution *p*(*Y_ijk_* | μ*_i_*, θ) = Poisson(*Y_ijk_* | λ*_ijk_*) if the conditional independence among detection events given λ*_ijk_*is assumed. Although we implemented an observation model for count detectors in this study, it can be easily extended to a model of varying effort with various detector types (e.g., binary proximity and multi-catch trap) as well as the basic SCR (Efford et al. 2013).

The population process of SCR describes the spatial pattern of activity centers following a point process in a region $ (Borchers and Efford 2008, Royle et al. 2009). $ is often defined as a rectangular region that includes an array of detectors. In this study, we applied a homogeneous Poisson process, the simplest form of the point process, assuming that μ follows a uniform distribution over $, μ ∼ Uniform($). Let ρ denote the log population density, and the number of activity centers in an arbitrary region with area *A* follows a Poisson distribution with expectation exp(ρ)*A*. As is the case with the basic SCRs (Borchers and Efford 2008), the model can be extended to an inhomogeneous Poisson process that can accommodate a situation in which ρ varies in space depending on the landscape covariates.

#### Home range formation

In ADCR, the utilization distribution is derived from a fine-scale movement behavior that forms a stationary home range in a heterogeneous permeability landscape. The Ornstein–Uhlenbeck process (Smouse et al. 2010) is a basic movement model describing the home range behavior of an animal that is subject to drift toward the activity center and a random walk. According to the circuit theory, permeability corresponds to the step length of a random walk (McRae et al. 2008), and we considered a situation in which permeability depends on landscape covariates. The Ornstein–Uhlenbeck process converges to a probability distribution of individual location at the equilibrium state (Smouse et al. 2010), and we applied it to utilization distribution in ADCR.

To derive the probability distribution of an individual location, we introduced an Eulerian approach that focuses on the flow of probability density in a space driven by drift and random walk. Under the Ornstein–Uhlenbeck process, the probability distribution of an individual location follows an advection-diffusion equation that describes the evolution of the probability density over time in space (Méndez et al. 2013). In our study, we consider a situation that the diffusion coefficient vary in space (Ovaskainen 2004, Ovaskainen et al. 2008) as follows:

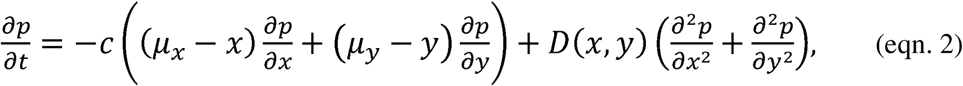

where *D*(*x*, *y*) > 0 is the spatially-varying diffusion coefficient and *c* is the intensity of drift. The first term on the right-hand side is advection corresponding to drift toward the activity center, and the second term is diffusion. The diffusion coefficient can be interpreted as mean step length of diffusive movement per unit time (Potts and Schlägel 2020) meaning how easy an animal can go through the location. In terms of circuit theory, it is equivalent to permeability or conductance (McRae 2006, McRae et al. 2008). The intensity of drift is the strength of site fidelity; the large value indicates that an animal is attracted toward the activity center strongly, resulting in small home range size. To model the effect of landscape variables on permeability, we applied the log-linear mode ln(*D*(*x*, *y*)) = β_0_ + β**z**(*x*, *y*), where β_0_ is the intercept, **z**(*x*, *y*) is a vector of the landscape covariates at location (*x*, *y*), and β is the corresponding regression coefficients that determine the dependence of permeability to landscape covariates. The intensity of drift, *c*, takes a positive value, and ln(*c*) is denoted by the parameter α_0_. To ensure that the total mass of probability does not change with time, the reflecting boundary condition is given for the space. Eqn. 2 describes the time evolution of the probability density of an individual location, and we applied it to project the animal movement paths for the data simulation (see **Simulation Study**).

Similar to mechanistic home range analysis (Moorcroft and Lewis 2006), the home range in this system is defined as the equilibrium probability density distribution *p*\*(**s** | μ), which emerges when advection toward activity center and diffusion balances. The *p*\*(**s** | μ) is obtained by solving the following system:

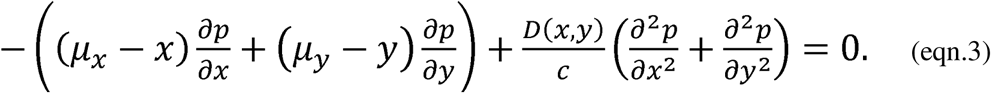

To make sure that *p*\*(**s** | μ) is the probability distribution in the region $, we imposed a constraint of ∫_c)_ *p* d**s** = 1. Note that the parameters α_0_ and β_0_ are redundant under the equilibrium assumption because ln(*D*(*x*, *y*)/*c*) = β_0_− α_0_ + β**z**(*x*, *y*). To ensure the identifiability of the model, it was reparametrized as γ_0_ = β_0_ − α_0_. γ_0_ is interpreted as the logarithm of the relative intensity of diffusion and drift, which determines the average home range size.

Although eqns. 2 and 3 imply that the landscape covariates are spatially continuous, they are actually given by a discrete dataset in raster or vector format. Accordingly, these equations can be numerically solved using a variety of discretization techniques, including the finite difference method and finite element method (Hirsch 2007). In this study, the finite difference method was applied because it is suitable for common situations in which landscape factors affecting the diffusion coefficient are available as a raster dataset. In addition, grid cells of raster data were used to discretize space for simplicity. Technical details for discretization and numerical derivation of *p*\*(**s** | μ) are shown in **Appendix S1**. Importantly, our home range specification is a generalization of basic SCRs that assume a bivariate normal home range (Borchers and Efford 2008, Royle et al. 2009), given that the solution of eqn. 3 when D(*x*, *y*) is constant over space (i.e., β = **0**) is a bivariate normal distribution whose variance is exp(γ_0_) (Méndez et al. 2013). An example of derived home ranges with different βs in a hypothetical landscape with a binary covariate for permeability is shown in Fig. 1, which shows that the larger the effect of a landscape covariate, the more the home range shape skews toward the habitat patch with high permeability.

**Fig. 1.**
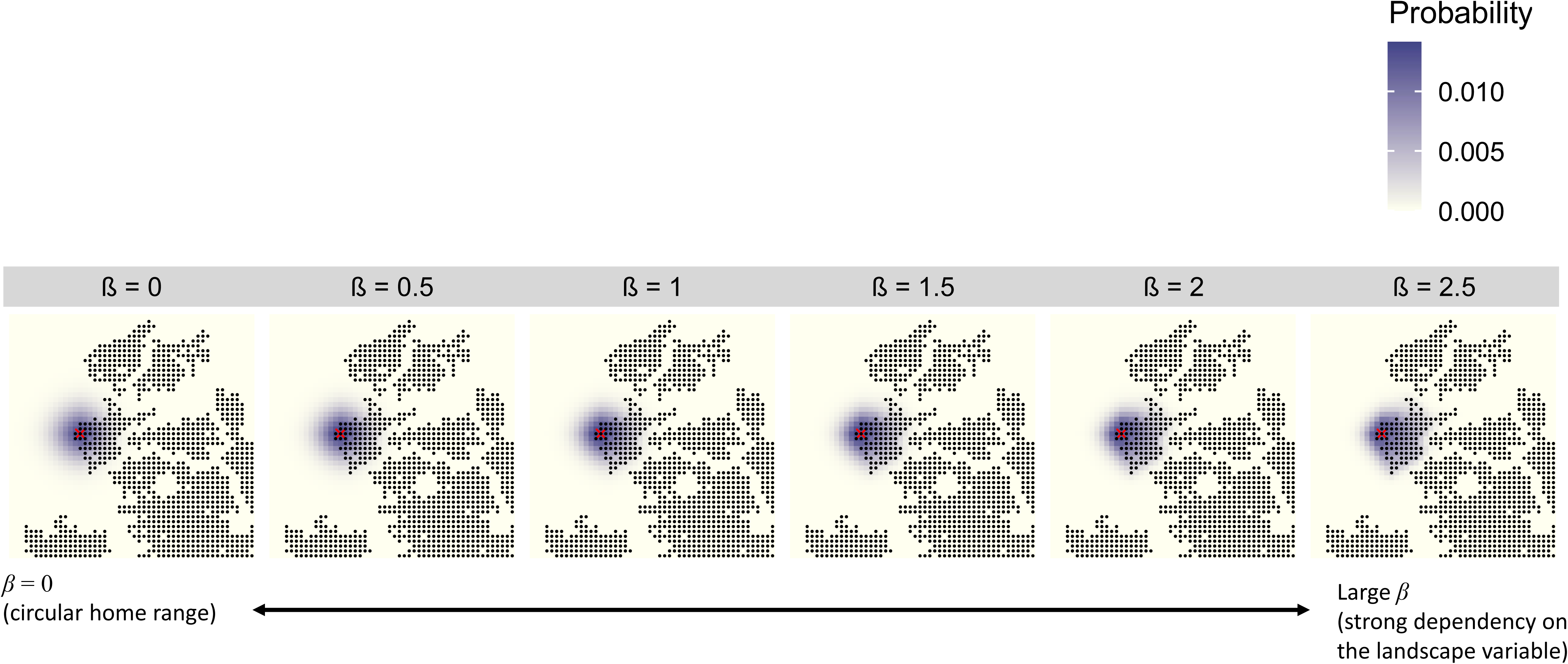
Example of simulated utility distributions under various dependencies of permeability on a hypothetical landscape variable. Black dots indicate cells with high permeability; other cells have low permeability. The red X indicates the activity center.

#### Model estimation

We applied the maximum likelihood method to estimate the permeability parameters and population density simultaneously from the capture–recapture data. Although conditional likelihood maximization can also be used for parameter estimation (Borchers and Efford 2008), full likelihood maximization was applied in this study because it has the capacity to extend to more complex situations in which population density varies across space. In this study, we applied a quasi-Newton algorithm (Nocedal and Wright 2006) for maximizing log likelihood function. Technical details in the model estimation was shown in **Appendix S2**.

#### Simulation study

Animal movement is temporally auto-correlated, and hence so are the locations of detections. To test the capacity of ADCR to estimate parameters of connectivity and population density correctly under a realistic movement process, we used a dataset generated via Monte Carlo simulations of animal movement. The landscape is a rectangular grid system of 50 × 50 (hence, the area is 2500), and animal location was described as a discrete coordinate of grid cells. Arrays of count detectors were set to the range, *x* = [9, 41] and *y* = [9, 41], with x and y axis spacing of 4 and 81 detectors. This simulation setting mimics a standard spatial capture recapture survey with a buffer outside the detector array to reduce boundary effects and with the distance between detectors so that an inidividual is detected by multiple detectors. The spatial resolution of the landscape grid was set coarsely to make the many iterative calculations in the simulation tractable.

Simulations were iterated 100 times, and true parameter values and landscape structure were randomly determined in each iteration. For each iteration, a heterogeneous landscape was generated using the R package rflsgen (Justeau-Allaire et al. 2022). Random patches were generated in the landscape by rflsgen with parameters shown in Table S1. For covariate values, 0.5 and −0.5 were assigned to patch and non-patch habitats, respectively. True parameter values were sampled from uniform distributions for each iteration. The ranges of true parameters were set as log population density ρ = (−3, −1), log intensity of drift α_0_ = (−1, 0) and coefficient of landscape variable β = (−2, 2). *g*_0_ and β_0_ were fixed to −3 and 1, respectively. At the beginning of an iteration, the number of individuals in the landscape was determined by a Poisson distribution with expected value of exp(ρ) × 2500 and activity centers were drawn from a discrete uniform distribution. The mean population size was 367 (range: 112, 956).

The animal movement simulation was run for *t* = 5000 after 100 burn-in steps for randomizing initial locations of individuals. For each time step, we calculated the probability vector of individual locations for all cells given the location at the previous step using the advection–diffusion equation with the explicit method (eqn. 4 in Appendix S1), and the location was sampled from a categorical distribution. When there was a detector in a cell in which an individual was located, the number of detection events was simulated by a Poisson distribution with expectation exp(*g*_0_). Then, single-occasion capture–recapture data were obtained by censoring undetected individuals. An example map of simulated movement and observations of an individual is shown in Fig. 2. The mean number of individuals detected at least once was 274 (range: 77, 682) with the detection rate (= number of detected individuals at least once / population size) of 0.745 (range: 0.631, 0.857). Mean number of detections was 2997 (range: 865, 7963). This simulation setting might generate a dataset with more detections than a real capture-recapture survey, but it will allow us to detect the bias of each estimator more sensitively.

**Fig. 2.**
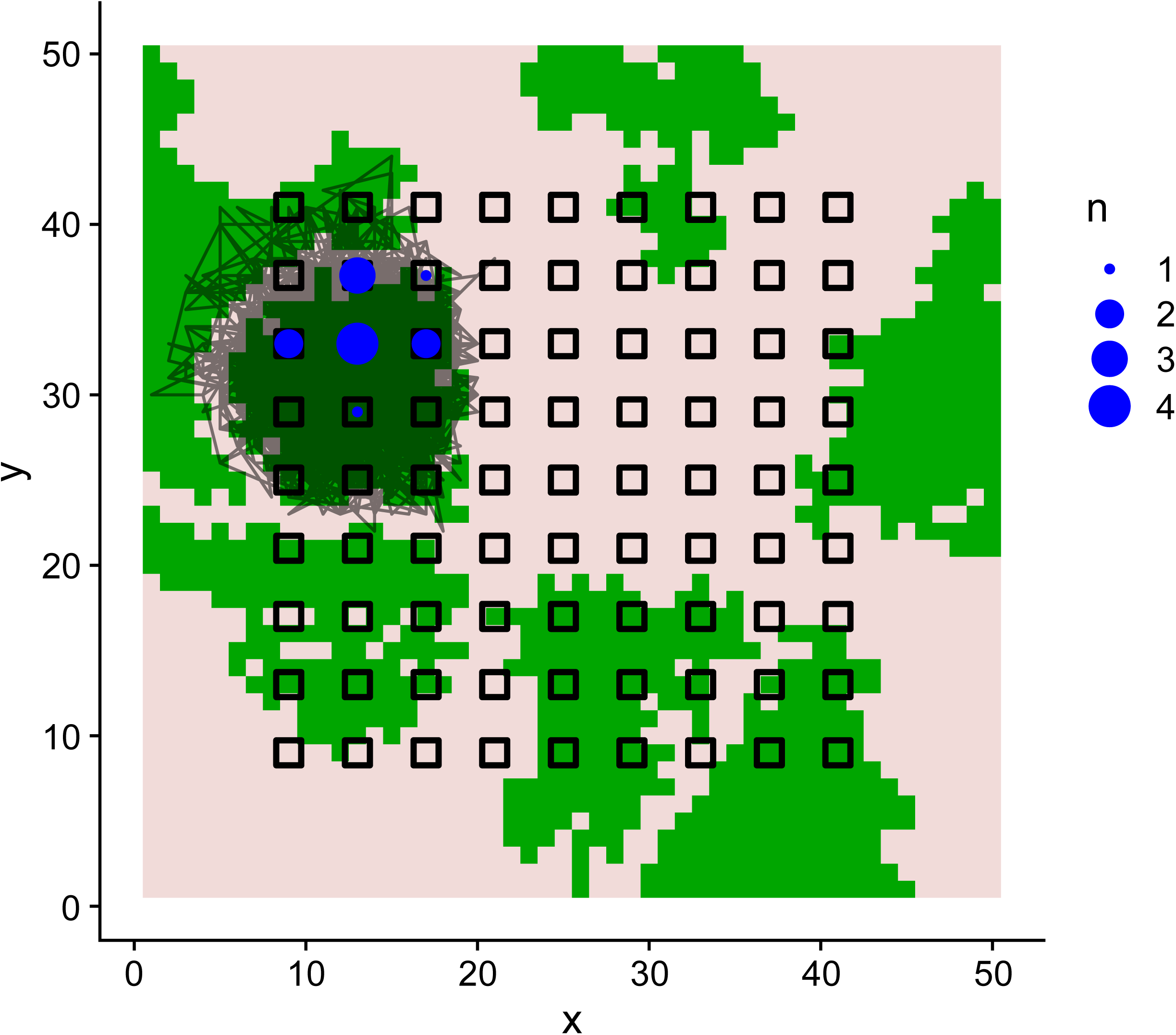
Example of simulated movement and the detection of an individual. The transparent solid line is the movement path. Open rectangles are traps, and blue circles indicate detection counts. The background tile map indicates zones with high (green) and low (pale red) permeability.

We applied ADCR, SCR with the least-cost path, and basic SCR to the simulated dataset to evaluate the accuracy of estimations. The basic SCR assumes that detectability function is Gaussian function of the Euclidian distance from the activity center ln(λ*_ijk_*) = g_0_ + *g*_1_||**s***_j_*- μ*_i_*||^2^, implying the home range shape is two-dimentional normal distribution. The formulation of SCR with the least-cost path is similar to basic SCR, but the Euclidian distance is replaced by the least-cost path distance from the activity center (*d_lcp_*(**s***_j_*,μ*_i;_ g*_2_)), ln(λ*_ijk_*) = g_0_ + *g*_1_*d_lcp_*(**s***_j_*,μ*_i;_ g*_2_). The log unit cost that an individual move from a grid cell (**s**) to the adjacent cell (**s’**) is *g*_2_(*z*(**s**) + *z*(**s’**)) / 2, and parameter *g*_2_ determines the weight of a landscape covariate on cost (Royle et al. 2013, Sutherland et al. 2015). Note that positive value of *g*_2_ indicates the negative relationship between the landscape connectivity and the landscape covariate, opposite to positive β value of ADCR.

For the three estimators, the deviation of population density estimates from true values was determined and root mean square error (RMSE) and mean bias error (MBE) were calculated. To show how well the three estimators reproduce home range shapes, the agreement between true (*p*\**_true_*) and predicted (*p*\* ) probability distributions of individual locations was calculated by an overlapping coefficient defined by the volume of overlapping region between two probability distributions (Inman and Bradley 1989). The overlapping coefficient ranges from 0 (no overlap) to 1 (perfect overlap). To evaluate the average performance over locations for activity center μ, we calculated the overlapping coefficients averaged over μ within the range of trap array x = [9, 41] and y = [9, 41] for each model and iteration. As an indicator of home range size, we also calculated 90% highest density areas of *p*\* whose μs are within the range of trap array and compared the means and SDs with the true values.

In order to demonstrate how well ADCR works under severe sparseness of observations and biased detector deployments to a specific landscape element, we conducted supplemental simulations with the following settings; true value of detectability was set to *g*_0_ = -5, and all the detectors in the landscape elements with low permeability (i.e. covariate value = -0.5) were removed from the detector array of the baseline simulation (range of detector quantities: 19 to 50). Other true parameters were chosen within the range of the baseline simulations: log population density ρ = -2, log intensity of drift α_0_ = -0.5, coefficient of landscape variable β = 2 and intercept of diffusion coefficient β_0_ = 1. Other settings, such as landscape structure, resolution, number of time steps were identical to the baseline simulations. The number of independent iterations was 100. The range of total population size was 296 to 386 (mean: 337). The number of individuals detected at least once ranged from 60 to 139 (mean: 96.8), and the detection rate was from 0.175 to 0.409 (mean: 0.287). The number of detections ranged from 90 to 230 (mean: 161).

All models were estimated using maximum likelihood methods with R 4.1.3 (R Core Team 2022). The movement simulation and log likelihood function of ADCR were written largely in R language, with some subroutines in C++. The source code is available online (https://github.com/kfukasawa37/adcrtest2). The log likelihood values of SCR with the least-cost path were derived based on the R script by Sutherland et al. (2015), with some modifications by Keita Fukasawa to deal with count detectors and the negative effect of landscape on the cost function. The basic SCR was estimated using the R package secr (Efford, 2022).

#### Case study

We used a capture-recapture dataset of Asiatic black bear population in eastern Toyama prefecture, Japan, collected over three years (2013-2015) by authors. The data was obtained using video-recording camera traps at 86 locations with a field design for efficient photographing of chest marks (Higashide et al. 2013) and photograph-based individual recognitions (Higashide et al. 2012). Note that all the camera locations were in forest due to limitations of field implementations. Years were treated as occasions. The dataset was constituted of 271 detections for 109 individuals with the locations and year. For the details of the capture-recapture survey, see **Appendix S3**.

In Japan, human-bear conflicts are increasing (Sakurai and Jacobson 2011). Understanding population status (Horino and Miura 2000) and space use (Ohnishi et al. 2019) of bears are crucial for developing plans to manage bear population and their threat to human well-being. Population and landscape genetic studies suggested that rivers (Saitoh et al. 2001, Ohnishi et al. 2007) and agricultural lands (Takahata et al. 2014, Ohnishi et al. 2019). To test the effect of rivers and agricultural lands as barriers or corridors for black bears, we compiled 0.5km grid data of area ratio covered by water surface and agricultural lands (Fig. S1). These areas were derived from the national vegetation map created by Biodiversity Center of Japan, Ministry of Environment (https://www.biodic.go.jp/index_e.html, accessed 24 May 2024) released in 2010s. We estimated ADCR and SCR with the least-cost path including the two landscape covariates and basic SCR. We compared the estimated log population density by the three methods, as well as the comparison of the estimated effect of landscape by ADCR and SCR with the least-cost path.

## Results

For all iterations, estimations of ADCR converged successfully. All the estimates of ADCR, including the population density and effect of landscape on connectivity, were substantially unbiased. The true and estimated parameters were on the line y = x, on average (Fig. 3). RMSE and MBE of *g*_0_ were 0.0258 and 0.0109, respectively. The true and predicted home ranges overlapped fairly well. The overlapping coefficients marginalized over activity centers were (0.950, 0.9996) over 100 iterations, and the mean was 0.988 (Fig. 4).

**Fig. 3.**
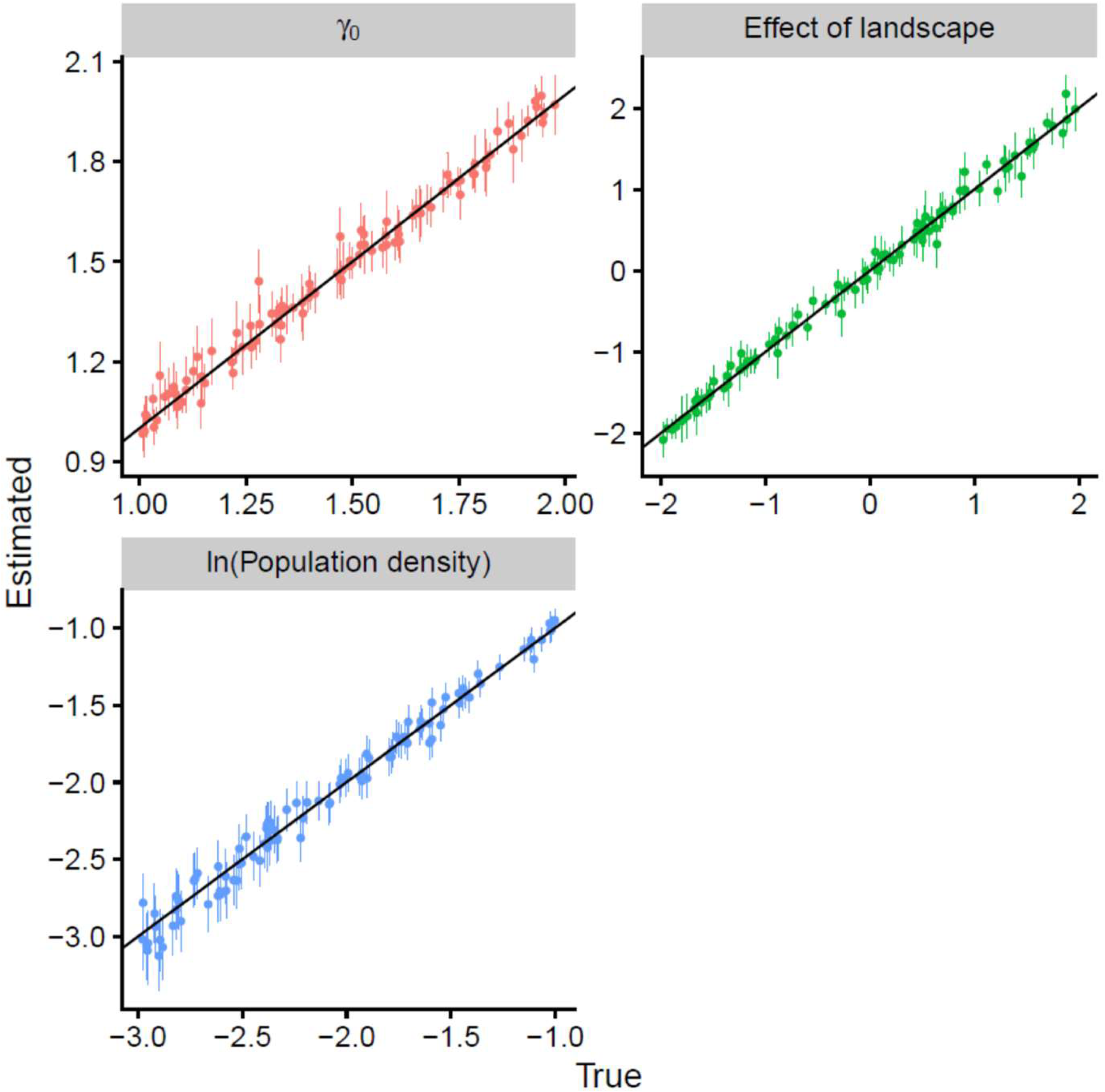
True vs. estimated parameters by ADCR. Dots and error bars indicate maximum likelihood estimates and 95% confidence intervals. The solid black line is y = x.

**Fig. 4.**
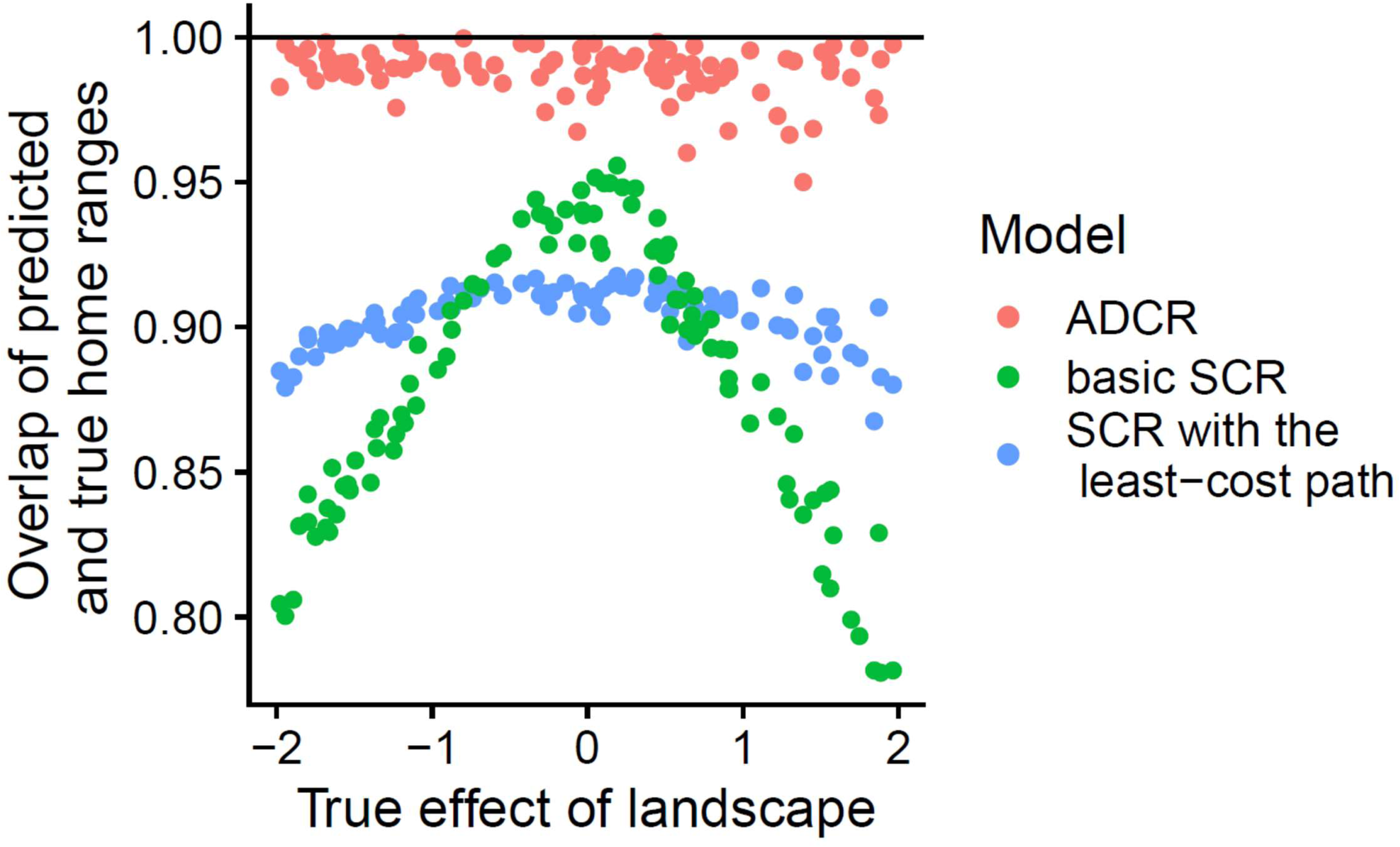
Overlap between true and predicted home ranges by ADCR, SCR with the least-cost path and basic SCR.

As predicted, population densities estimated by SCR with the least-cost path and basic SCR were robust to true connectivity settings in our simulation experiments. RMSEs and bias of log population density estimated by SCR with the least-cost path and basic SCR were very similar to those of ADCR and were not sensitive to the true effect of landscape on connectivity (Table 1). The coefficient of landscape estimated by SCR with the least-cost path were linear with the true parameters of connectivity, on average, but showed a slightly weaker relationship (Pearson’s *r* = −0.988, Fig. S2) than that of ADCR (Pearson’s *r* = 0.995, Fig. 3). The estimates of γ_0_ by ADCR were also linear with the true values (Pearson’s *r* = 0.991, Fig. 3). Although the mean home range sizes measured by 90% highest density area were almost unbiased for the three estimators (Fig. S3a), the heterogeneity in home range sizes were largely underestimated by SCR with the least-cost path and basic SCR (Fig. S3b); MBE in standard deviation of home range sizes were -5.38 for SCR with the least-cost path, -15.3 for basic SCR and -0.0508 for ADCR.The overlapping coefficients of home range predicted by SCR with the least-cost path (mean: 0.904; variance: 9.93×10^-5^) and basic SCR (mean: 0.883; variance: 2.23×10^-3^) were lower than those for ADCR (mean: 0.998; variance: 7.52×10^-5^) over the range of true connectivity parameters. The predictive performance of SCR with the least-cost path and basic SCR were decreased when the connectivity strongly depends on landscape covariate, but SCR with the least-cost path worked better than basic SCR (Fig. 4). For example of the range where the absolute value of true landscape effect is [1.5, 2.0], the mean overlaps by SCR with the least-cost path, basic SCR and ADCR were 0.892, 0.822 and 0.991, respectively. SCR with the least-cost path tended to predict a more blurred and isotropic home range shape than the true home range shape (Fig. S4).

**Table 1.**
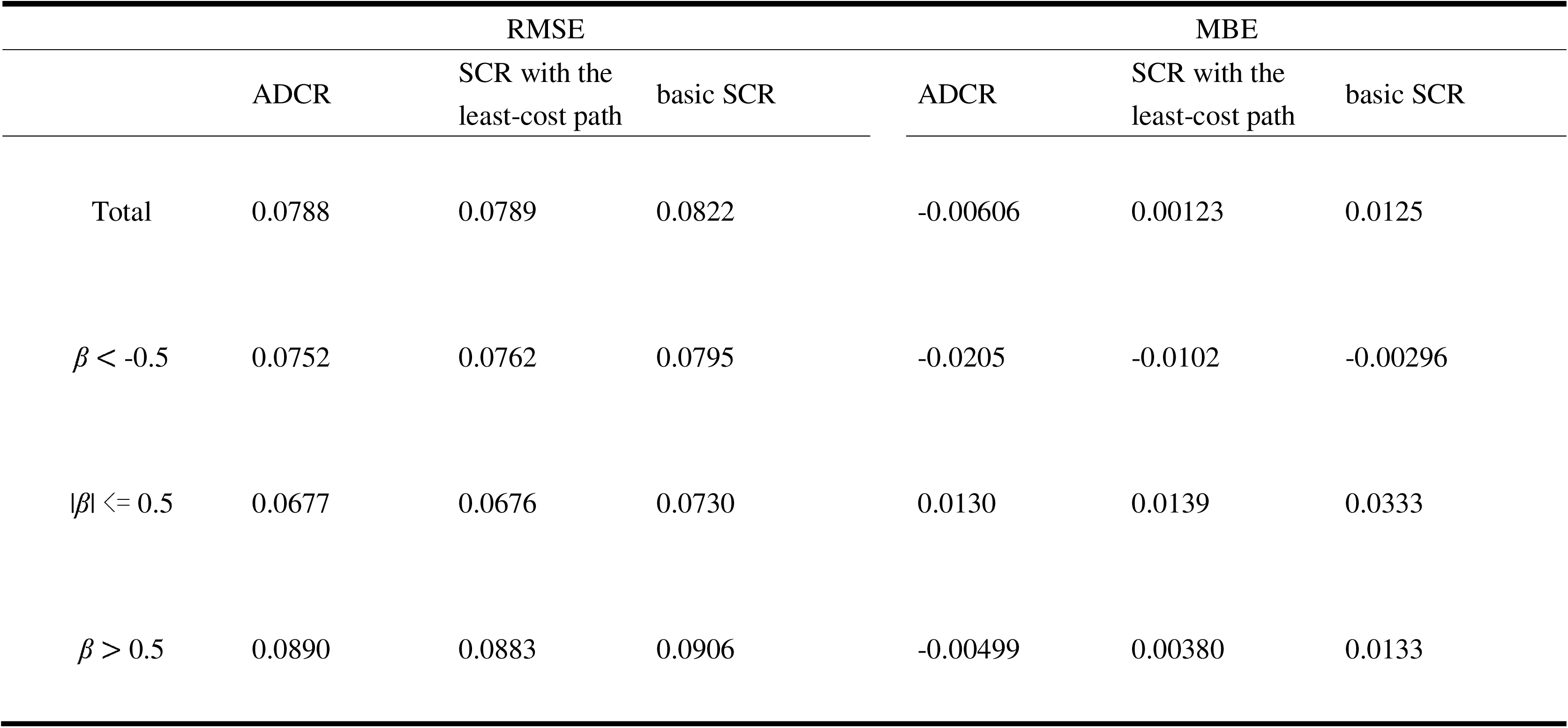
Root mean squared error (RMSE) and mean bias error (MBE) of log density for the three estimators for the baseline scenario. The values binned into three classes of the true landscape effect, β, were also shown.

Under the sceanio of severe data sparseness and biased detector alignment, estimates of ADCR had large variances but were substantially unbiased; the mean ± standard deviation of estimated log population density, γ_0_ and effect of landscape were -2.00±0.142, 1.41±0.425 and 2.12±1.10, respectively. In this scenario, accuracy of density estimates by ADCR was much better than the other estimators (Fig. S5); the MBEs of population density estimates by ADCR, SCR with the least cost path and basic SCR were 0.00308, 0.0657 and -0.0373, respectively. ADCR could detect the correct environmental signals on connectivity clearer than SCR with the least-cost path; the 95% confidence intervals (CIs) did not overlap 0 for 60 out of 100 iterations which is outperformed SCR with the least-cost path (54 out of 100 iterations). Although the sign (i.e. positive or negative) of landscape effect on connectivity was incorrect for two iterations of ADCR and five iterations of SCR with the least-cost path out of 100 iterations, all of them were not significant with their 95% CI overlapping 0 (Table S2). There were two iterations that SCR with the least-cost path estimated incorrect signs of effects and ADCR could detect significant effects correctly.

For the applications on the black bear dataset, there were unexpected discrepancies in the estimated effects of landscape factors on connectivity between ADCR and SCR with the least-cost path (Fig. 5). The estimated coefficients (i.e. parameter β) of agricultural land and water surface on the permeability of black bears were 0.361 (SE: 0.113; 95% CI: 0.175, 0.547) and -6.41 (SE: 1.26; 95% CI -8.88, -3.94), respectively. Inversely, the estimated effect of water surface on the cost by SCR with the least-cost path (parameter *g*_2_) was -25.5 (SE: 5.13; 95% CI: -35.6, -15.4), indicating water surface was corridor for black bears (Fig. 5). Note that deletion of agricultural land from the covariate did not affect the effect of water surface so much (-23.1, SE: 4.57). The effect of agricultural land by SCR with the least-cost path was not clear; the maximum likelihood estimate was 0.0692 (SE: 0.0360; 95% CI: -0.00130, 0.140) (Fig. 5). Contrary to the predicted home ranges by ADCR showing skewed shapes avoiding water surface (Fig. S6a), the home ranges predicted by SCR with the least-cost path showed “leakage” of probability density into water surface (Fig. S6b). The log population density estimates were similar among the methods but ADCR was the highest: -1.45 (SE: 0.113; 95% CI: -1.67, -1.23) by ADCR, -1.56 (SE: 0.0996; 95% CI: -1.76, -1.37) by SCR with the least-cost path and -1.59 (SE: 0.105; 95% CI: -1.79, -1.37) by basic SCR.

**Fig. 5.**
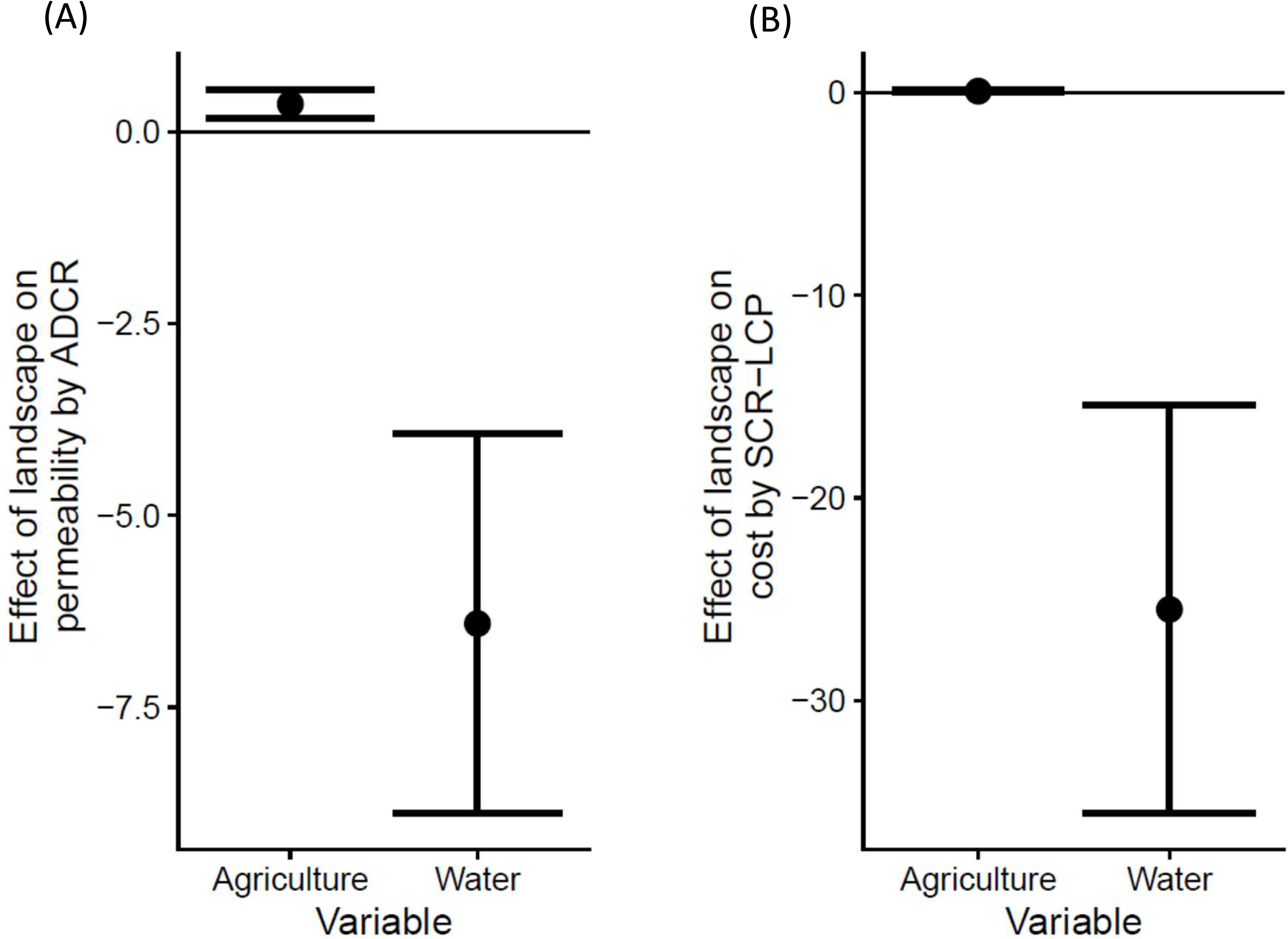
Estimated effect of landscape on (A) permeability by ADCR and (B) cost by SCR with the least-cost path (SCR-LCP) for black bear dataset. Dots and error bars are the maximum likelihood estimate and 95% CIs, respectively.

## Discussion

Individual movement is an important driver of spatial patterns of populations, and incorporating movement processes in spatial capture–recapture analysis will facilitate studies of the interplay between individual- and population-level ecological processes (Royle et al. 2018, McClintock et al. 2022). In this study, we developed a formal framework for a SCR model including mechanistic home range formation with an explicit theoretical link to animal movement models. Through individual-based simulations of animal movement and detection, we showed ADCR accurately estimated the population density, effect of landscape on permeability and heterogeneous home range sizes. For the application of models to black bear dataset, ADCR could detect biologically-plausible environmental signals on the connectivity. These results suggest that ADCR is an effective approach for estimating population density and random walk-based landscape connectivity.

Although the integration of a least-cost path modeling with spatial capture recapture analysis (Royle et al. 2013, Sutherland et al. 2015) was a great methodological advancement for the simultaneous estimation of population density and ecological distance, the model performance under the animal movement following step selection function has not been studied well. Our simulation study demonstrated that ADCR outperformed the SCR with the least cost path in terms of accracy in population density estimation, successful detection of environmental signals on connectivity (i.e. permeability or cost) and precise prediction of home range shapes and the location-specific heterogeneity in home range sizes. The ability of ADCR to estimate the effects of landscape variable on animal movement correctly (Fig. 3) is result from the formal link between microscopic animal movement and the macroscopic utility distribution, corresponding to Lagrangian and Eulerian descriptions of a system. Previous study showed that SCR with the least-cost path is difficult to estimate accurate cost on movement without an integration with GPS logging data (Dupont et al. 2022). This might be due to lack of formal link between home range model and the underlying movement processes.

For the case study of Asiatic black bears, the results of ADCR supported the previous studies demonstrated rivers as physical barriers (Saitoh et al. 2001, Ohnishi et al. 2007) and farmlands as corridors (Takahata et al. 2014), whereas SCR with the least-cost path showed the unexpected estimates indicating water surface as corridor for bears. Although our simulation study with sparse data and biased detector alignment could not fully reproduce such an extraordinary discrepancy between estimators, there were some cases that only ADCR could detect correct environmental signals on connectivity (Table S2). We considered the unexpected behavior of SCR with the least-cost path was estimation bias caused by combination of biased layout of detectors toward forest and the model assumption that detection rate at the activity center is constant irrespective of the home range size. This assumption implys that observations do not reflect the ubiquitous tradeoff between exploration of large area and exploitation of resources near the activity center (exploration-exploitation tradeoff, Eliassen et al. 2007, Berger-Tal et al. 2014, Berger-Tal and Saltz 2014), leading model vulnerability in estimating of animal space use to sampling non-randomness. ADCR explicitly models home ranges as probability distributions with sum-to-one constraint, and seemed to be more suitable for general home range behavior. However, it may not be able to fully handle a variety of the diverse animal movement processes in response to landscape barriers such as the correlated random walk (Patlak 1953, Wang and Potts 2017) and randomized shortest paths (Panzacchi et al. 2016, Long 2019). Extending the current study, which integrated the utility distribution derived from the stochastic process of animal movement with spatial capture-recapture models, to a variety of movement models will lead to solve the problem.

The effect of a skewed home range on the accuracy of density estimation by SCR has been studied extensively in ecology (Ivan et al. 2013, Royle et al. 2013, Sutherland et al. 2015, Efford, 2019, Theng et al. 2022). The results of this study were consistent with those of recent studies (Efford 2019, Theng et al. 2022) showing that robustness of density estimation to deviations in assumptions on home range shapes depends on independence of sampling design to animal space use. In addition, our study showed that estimation bias of density by SCR with the least-cost path can be as large as basic SCR (Fig. S5). Efford (2019) emphasized the importance of sampling design in space, such as the use of a two-dimensional detector array and avoidance of detector array alignment along landscape features, for the correct estimation of density. Obviously, such sampling designs are also favorable for successful detection of landscape effects on home range shape by ADCR.

In landscape ecological analyses, effects of choice of spatial resolution on the results of study can often be problematic (Turner et al. 1989, Qi and Wu 1996). We found that ADCR was robust to decreased spatial resolution of a landscape variable by supplementary simulatios (Appendix S4, Fig. S7). This robustness is a favorable characteristic of connectivity measures in circuit theory, which shares the same theoretical foundation of random walk descriptions of animal movement (Hanks and Hooten 2013).

The broad application of SCR in ecology can largely be attributed to its flexibility to incorporate heterogeneity in ecological processes underlying spatial capture recapture data, such as environment-dependent abundance and sex-biased detection (Efford, 2022). Future advances in ADCR should focus on similar functionality. In particular, incorporating heterogeneity in home range size (Efford et al., 2016) as well as heterogeneity in connectivity is a major challenge. Also, the generalization of ADCR for dealing with temporally open populations driven by birth-death processes (Gardner et al. 2010, Efford & Schofield 2022) and integration with GPS logging data (Dupont et al. 2022) would be helpful for getting more out of capture–recapture data. The development of software tools that enable users to benefit from the flexible nature of ADCR is an urgent task for expanding the application of the method.

Spatial capture–recapture is a popular method for abundance analyses in ecology and conservation (Iijima 2020), and large amounts of data have accumulated. Moreover, recent technical advances in individual identification from environmental DNA (Dugal et al. 2022) and machine observations, such as camera traps (Schneider et al. 2022), will expand the applicability of capture–recapture approaches to broad taxa. Such capture–recapture data have the potential to advance our understanding of the process of animal home range formation. ADCR will be a useful tool for integrated analyses of capture recapture data incorporating individual- and population-level ecological processes.

## Acknowledgements

Yutaka Osada provided valuable comments on an earlier version of the statistical model. We thank Elise Zipkin and two anonymous reviewers for constructive comments in the peer-review process. This study was supported by JSPS KAKENHI Grant Numbers JP13247736, JP19153418, and by the Environment Research and Technology Development Fund (JPMEERF20194005) of the Environmental Restoration and Conservation Agency Provided by the Ministry of Environment of Japan.

## Author contributions

KF conceived the ideas, designed the methodology, conducted the simulation study, conducted field survey, and applied the model to actual dataset. DH conducted field survey, and complied the dataset. KF led the writing and all authors contributed to the revision of the manuscript.

## Supporting information

**Appendix S1** Technical details in the finite difference method.

**Appendix S2** Technical details in model estimation with maximum likelihood methods.

**Appendix S3** Detailed descriptions on capture-recapture survey data used for case study.

**Appendix S4** The effect of spatial resolution on parameter estimates.

**Appendix S5** Supplementary figures.

Fig. S1 (a) Study area, and (b) locations of camera traps (red rectangle), agricultural lands (orange) and water surface (blue), and 0.5km grid cells.

Fig. S2 Relationship between the true effect of landscape on permeability and estimated effect of landscape on cost by SCR with the least-cost path. Error bars are 95% CIs.

Fig. S3 Summary statistics of 90% highest density (HD) area of true and predicted home ranges, (a) means and (b) standard deviations (SDs). A dot corresponds to an iteration of the simulation, and the means and SDs are among home ranges with different activity center locations.

Fig. S4 Example of true and predicted home ranges by ADCR and SCR with the least-cost path. Red x symbols mark the home range center. Black dots are zones with high permeability.

Fig. S5 Estimated log population density by ADCR, SCR with the least-cost path (SCR-LCP) and basic SCR under the scenario of severe data sparseness (i.e. g0 = -5) and biased detector alignment. Black dots are maximum likelihood estimates (MLEs). Open circles and and error bars indicate the mean and SD of MLEs. Horizontal black lines are the true value (-2.0).

Fig. S6 Examples of predicted home ranges by (a) ADCR and (b) SCR with the least-cost path for the black bear. Red x symbols mark the home range centers. Black dots are cells contain water surface.

Fig. S7 Comparison between the estimates of baseline simulations and 1/2 resolution scenarios for (a) effects of landscape and (b) log population densities by ADCR. Dots and error bars are maximum likelihood estimates and 95% CI, respectively. Glay lines are principal components of parameter estimates.

**Appendix S6** Supplementary table.

Table S1 Target parameter ranges for the neutral landscape generator, rflsgen. The parameter names were based on Justeau-Allaire et al. (2022). Parameters not shown were set to default values.

Table S2 Contingency table on the correctness of sign (i.e. positive or negative) of landscape effect on connectivity estimated by ADCR and SCR with the least-cost path for the simulation scenario of sparse data and biased detector alignment. The significance was determined by non-overlap of 95% confidence interval with 0.

