## Appendix S1 for "Mechanistic home range capture–recapture models for the estimation of population density and landscape connectivity"

**Appendix S1** Technical details in the finite difference method.

We applied a finite difference method to derive a numerical solution to an advection–diffusion model of utilization distribution in a discretized space. In this study, a finite difference method was applied in dynamic setting for the animal movement simulation as well as static setting for estimation of ADCR.

Discretization schemes for a finite difference method have trade-offs between precision and numerical stability (Hirsch 2007). To ensure numerical stability for a diverse range of parameter values, we applied a first-order upwind scheme for advection and second-order central difference scheme for diffusion. Discretizing time by backward differentiation, eqn. 2 is approximated by a transition matrix for a cell-wise probability vector **p** = (*p*_1_, *p*_2_, …, *p_N_*) is obtained as follows:
$\mathbf{p}_{t+\Delta t}=\left\{ \mathbf{I}+\left( -c\left( \frac{\mathbf{A}_{x1}\mathrm{diag}\left( \left| \mu_{x}-\mathbf{x} \right| \right)}{\Delta x}+\frac{\mathbf{A}_{y1}\mathrm{diag}\left( \left| \mu_{y}-\mathbf{y} \right| \right)}{\Delta y} \right)+\mathrm{diag}\left( \mathbf{d}^{1/2} \right)\left( \frac{\mathbf{A}_{x2}}{{\Delta x}^{2}}+\frac{\mathbf{A}_{y2}}{{\Delta y}^{2}} \right)\mathrm{diag}\left( \mathbf{d}^{1/2} \right) \right)\Delta t \right\}\mathbf{p}_{t}$	(eqn. 4)
where **A***_x_*_1_ and **A***_y_*_1_ are non-symmetric adjacency matrices corresponding to the advection term for the *x* and *y* axis, respectively. **A***_x_*_2_ and **A***_y_*_2_ are symmetric adjacency matrices corresponding to the diffusion term. Off-diagonal elements (*i,* *j*, *i≠j*) in the adjacency matrix for advection are 1 if cell *i* and *j* are neighbors and the direction of drift is *i* -> *j* and are 0 otherwise. Off-diagonal elements in the adjacency matrix for diffusion are 1 if cell *i* and *j* are neighbors. To satisfy the principle of probability mass conservation, diagonal elements of both types of adjacency matrices were determined so that all column sums are 0. Δ*x*, Δ*y* and Δ*t* are resolutions of discretization for the *x* axis, *y* axis, and time, respectively. **x**, **y**, and **d** are vectors of the *x* and *y* coordinates and diffusion coefficient of grid cells, respectively. The function diag() indicates vector-to-diagonal matrix conversion. The diffusion coefficient **d** = (*d*_1_, *d*_2_, …, *d_N_*) is positive-valued and often assumed to be dependent on cell environmental variables (e.g., topographic and land cover variables).

Using an explicit method, **p***_t_*_+Δ_*_t_* is projected from **p***_t_*. A realization of animal movement at time *t*+Δ*t* follows categorical distribution with probability vector **p***_t_*_+Δ_*_t_*. This is a discrete space analogue of the step selection function defined by the advection-diffusion equation (Potts and Schlägel 2020), and this stochastic model was used to simulate animal movement.

Home range in this system is defined as an equilibrium utility distribution, *p**, which is the solution of *p* when δ*p*/δ*t* = 0. Discrete approximation of the utility distribution, **p***, is given by solving the following sparse linear system:
$\left( -\left( \frac{\mathbf{A}_{x1}\mathrm{diag}\left( \left| \mu_{x}-\mathbf{x} \right| \right)}{\Delta x}+\frac{\mathbf{A}_{y1}\mathrm{diag}\left( \left| \mu_{y}-\mathbf{y} \right| \right)}{\Delta y} \right)+\mathrm{diag}\left( \left( \frac{\mathbf{d}}{c} \right)^{\frac{1}{2}} \right)\left( \frac{\mathbf{A}_{x2}}{{\Delta x}^{2}}+\frac{\mathbf{A}_{y2}}{{\Delta y}^{2}} \right)\mathrm{diag}\left( \left( \frac{\mathbf{d}}{c} \right)^{\frac{1}{2}} \right) \right)\mathbf{p}=\mathbf{0}$.
(eqn. 5)

Given *μ*_x_, *μ_y_* and **d**/*c*, this equation has a unique solution with constraint **1**^T^**p =** 1.
