## Appendix S2 for "Mechanistic home range capture–recapture models for the estimation of population density and landscape connectivity"

**Appendix S2** Technical details in model estimation with maximum likelihood methods.

As is the case with the basic SCRs (Borchers and Efford 2008), the likelihood that ADCR will be formulated by a product of the Poisson process likelihood that *n* individuals were observed at least once and the conditional likelihood of capture history **ω** = (**ω**_1_**, ω**_2_, ..., **ω***_n_*) marginalized over activity centers given *n* individuals. Let $p_{.}\left( \boldsymbol{\mu,\theta} \right)$ denote the probability that an individual whose activity center is **μ** is detected at least once, $p.\left( \boldsymbol{\mu},\boldsymbol{\theta} \right)=1-\prod_{j} \prod_{k} p\left( Y_{jk}=0|\boldsymbol{\mu},\boldsymbol{\theta} \right)$, and $p\left( \boldsymbol{\omega}_{i}|\boldsymbol{\mu}_{i},\boldsymbol{\theta} \right)$ denote the unconditional probability of capture history, $p\left( \boldsymbol{\omega}_{i}|\boldsymbol{\mu}_{i},\boldsymbol{\theta} \right)=\prod_{j} \prod_{k} p\left( Y_{ijk}|\boldsymbol{\mu}_{i},\boldsymbol{\theta} \right)$. Then, the full likelihood is as
$L\left( \boldsymbol{\theta},\rho\right)=\frac{\left\{ exp\left( \rho\right)\int_{\mathcal{S}} p_{.}\left( \boldsymbol{\mu,\theta} \right)d\boldsymbol{\mu} \right\}^{n}exp\left( -exp\left( \rho\right)\int_{\mathcal{S}} p_{.}\left( \boldsymbol{\mu,\theta} \right)d\boldsymbol{\mu} \right)}{n!}\times\left( \begin{matrix} n \\ n_{1},n_{2},\ldots,n_{C} \end{matrix} \right)\prod_{i=1}^{n} \frac{\int_{\mathcal{S}} Pr\left( \omega_{i}|\boldsymbol{\mu},\boldsymbol{\theta} \right)d\boldsymbol{\mu}}{\int_{\mathcal{S}} p_{.}\left( \boldsymbol{\mu,\theta} \right)d\boldsymbol{\mu}}$ ,(eqn. 6)
where *n*_1_, *n*_2_, …, *n*_C_ are the frequencies of unique capture histories. Although conditional likelihood maximization can also be used for parameter estimation (Borchers and Efford 2008), full likelihood maximization was applied in this study because it has the capacity to extend to more complex situations in which population density varies across space. Automatic differentiation of likelihood functions (Auger-Méthé et al. 2017) is not currently available for ADCR, but a quasi-Newton algorithm, such as the Broyden–Fletcher–Goldfarb–Shanno (BFGS, Nocedal and Wright 2006) algorithm with numerical differentiation of log likelihoods, works well.
