## Appendix S3 for "Mechanistic home range capture–recapture models for the estimation of population density and landscape connectivity"

**Appendix S3** Detailed descriptions on capture-recapture survey of Asiatic black bears

### Study site

Our survey was conducted in the eastern Toyama prefecture, Japan. Our study site locates at the western foot of Tateyama mountains and partly overlapped to the Chubusangaku National Park. It contains a wide range of topography from lowland, hill to mountains. In the hilly area, agricultural lands along the rivers divide the forest landscape. The deciduous coniferous trees (*Fagus crenata*, *Quercus crispula* and *Q. serrata*) which offer food for bears in autumn are dominant species of the forest (Arimoto et al. 2011). As in other parts of Japan, a poor crop of beechnuts and acorns causes behavioral changes in black bears that increase conflicts with human (Ohnishi et al. 2011).

### Survey design

From 2013 to 2015, we conducted a camera trap capture-recapture survey at 86 locations (Fig. S1). The survey were conducted from June to October, which is active season for bears. In each location, we set a camera trap (Trophycam 119437C; Bushnell Outdoor Products, Overland Park) with video-recording mode. The duration of video was 30 seconds, and lag time after a trigger was set to 10 seconds. For efficient photographing of a chest mark as a key to individual recognition, we used an odor stimulant (mixture of honey and red wine) to encourage bears to stand up in front of camera by the protocol shown by Higashide et al. (2013). In this protocol, a spruce-pine-fir (SPF) lumber was fixed horizontally at a height of 1.5 m on two tree trunks in each camera trap’s field of view as a grub bar for bears to stand up. Then, the odor stimulant was filled in a plastic bottle covered by a robust polyvinyl chloride tubing for protection from bear attacks and fixed to the center of SPF lumber. Note that all of the our camera trap locations are in forest because of the limitations in field implementation. We visited each location every one to two months to replace batteries and SD cards and to refill the odor stimulant. The records of the same individual at a location within 30 minutes were grouped into a detection event. An image library of chest marks was developed from the video footage taken, and identical individuals were matched manually (Higashide et al. 2012). For fitting the capture recapture models, we aggregated the numbers of detection events for each camera trap, individual and year.
