## Appendix S4 for "Mechanistic home range capture–recapture models for the estimation of population density and landscape connectivity"

**Appendix S4** The effect of spatial resolution on parameter estimates

The spatial resolution of landscapes can often alter the results of landscape ecology, and the robustness of models to differences in spatial resolution between data generation and model estimation should be evaluated. To evaluate the robustness of ADCR to a mismatch in spatial resolution of landscape data with the resolution that matters to the actual movement of animals, I considered an additional scenario of the simulation study in which the available landscape variable was restricted to 1/2 resolution. For the 1/2 resolution scenario, grid cells with side length 2 were used for discretizing space in the model estimations, and the landscape variable was averaged for each grid cell. Simulated detection matrices and methods for model estimation were same as the baseline simulation. I compared the density and connectivity estimates for the 1/2 resolution scenario with estimates for the baseline scenario.

ADCR was robust to mismatches in landscape resolution between the data generating process and model estimation. The effect of landscape and log population density in the 1/2 resolution scenario were almost the same as those for the baseline scenario (Fig. S7, slopes of principal axes of the effect of landscape and log population density were 1.05 and 0.999, respectively).
