## Appendix S5 for "Mechanistic home range capture–recapture models for the estimation of population density and landscape connectivity"

Appendix S5 Supplementary Figures.

Fig. S1 (a) Study area, and (b) locations of camera traps (red rectangle), agricultural lands (orange) and water surface (blue), and 0.5km grid cells.

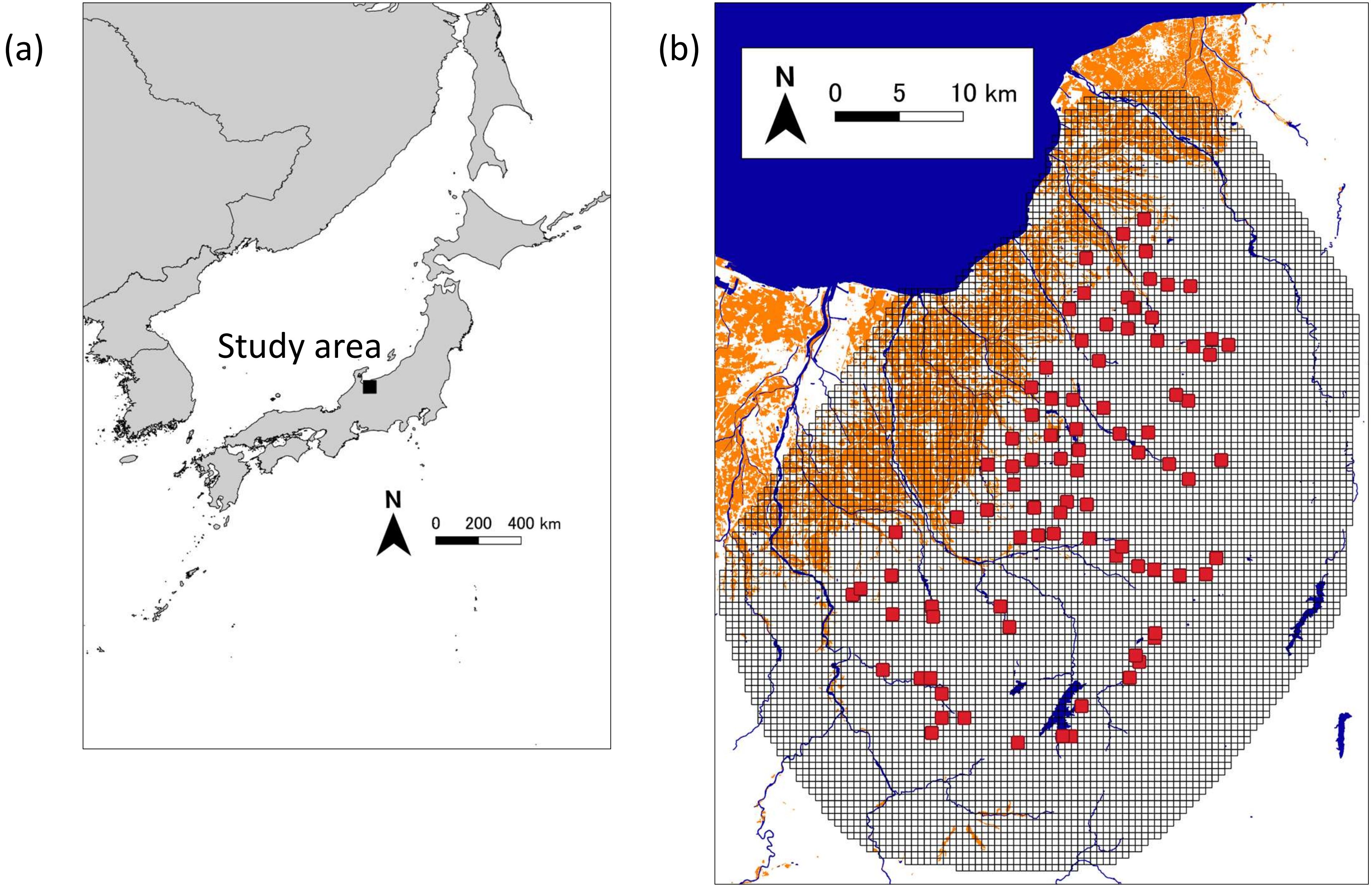

Fig. S2 Relationship between the true effect of landscape on permeability and estimated effect of landscape on cost by SCR with the least-cost path (SCR-LCP). Error bars are 95% CIs.

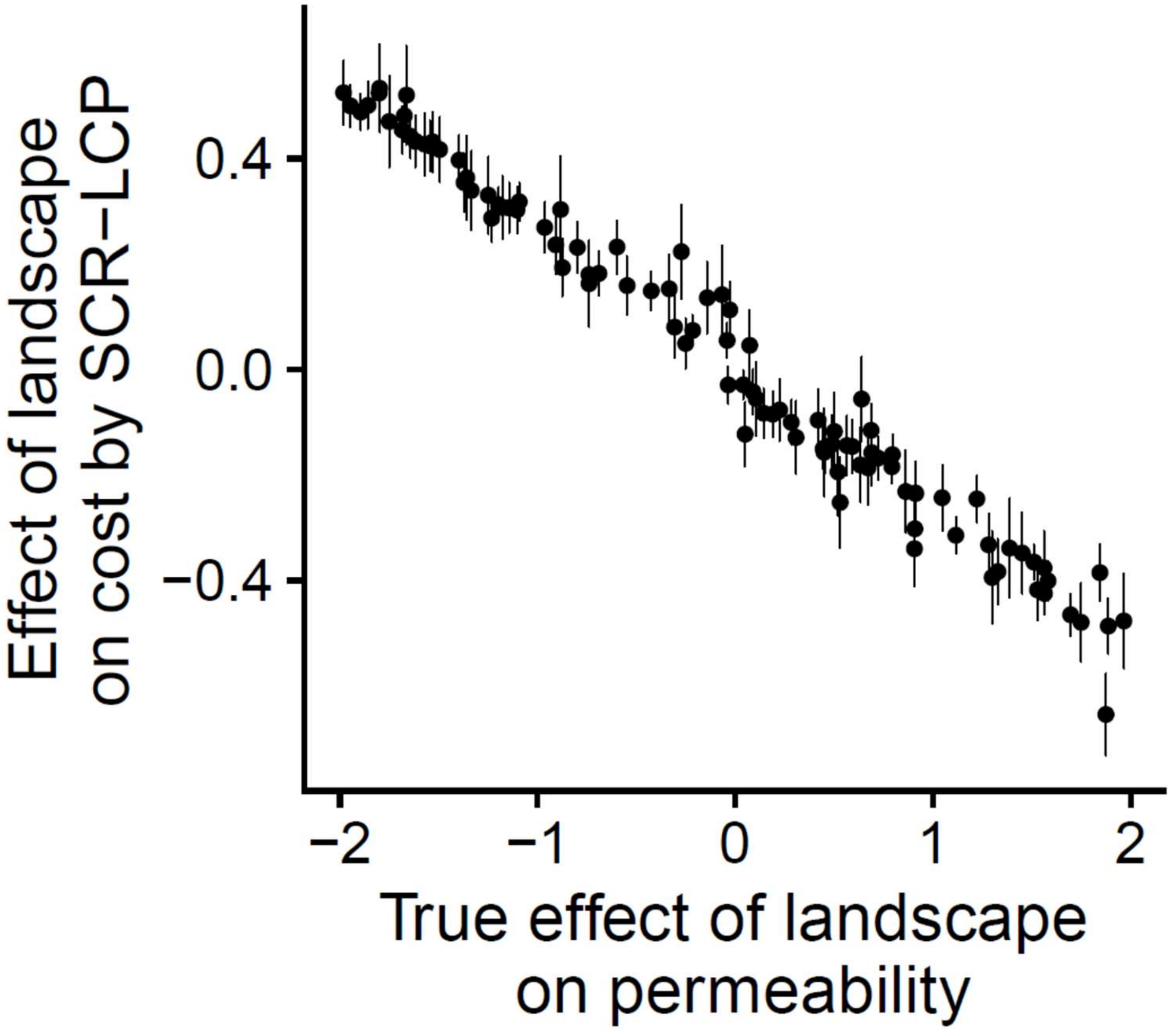

Fig. S3 Summary statistics of 90% highest density (HD) area of true and predicted home ranges, (a) means and (b) standard deviations (SDs). A dot corresponds to an iteration of the simulation, and the means and SDs are among home ranges with different activity center locations.

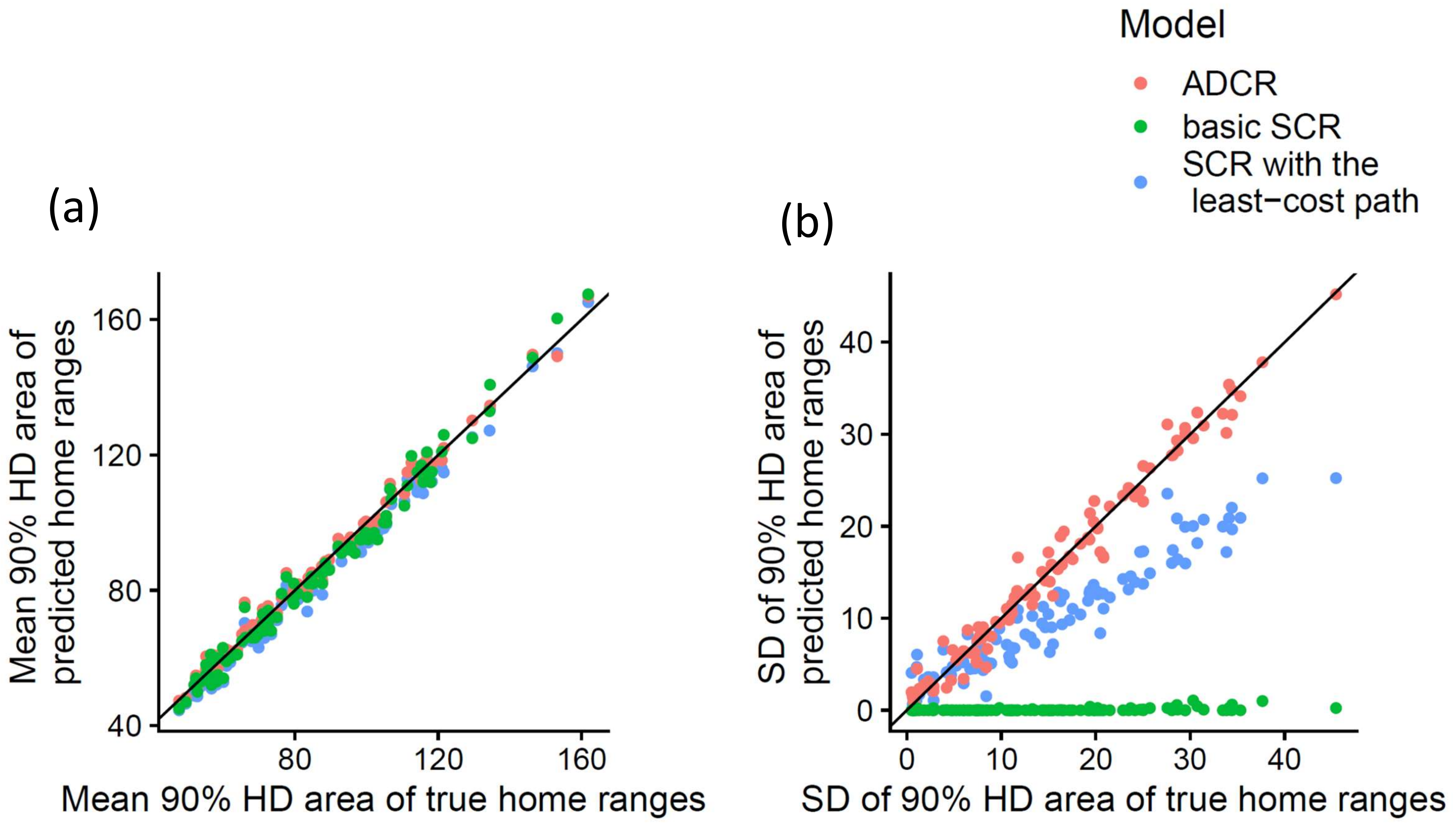

Fig. S4 Example of true and predicted home ranges by ADCR and SCR with the least-cost path. Red x symbols mark the home range center. Black dots are zones with high permeability.

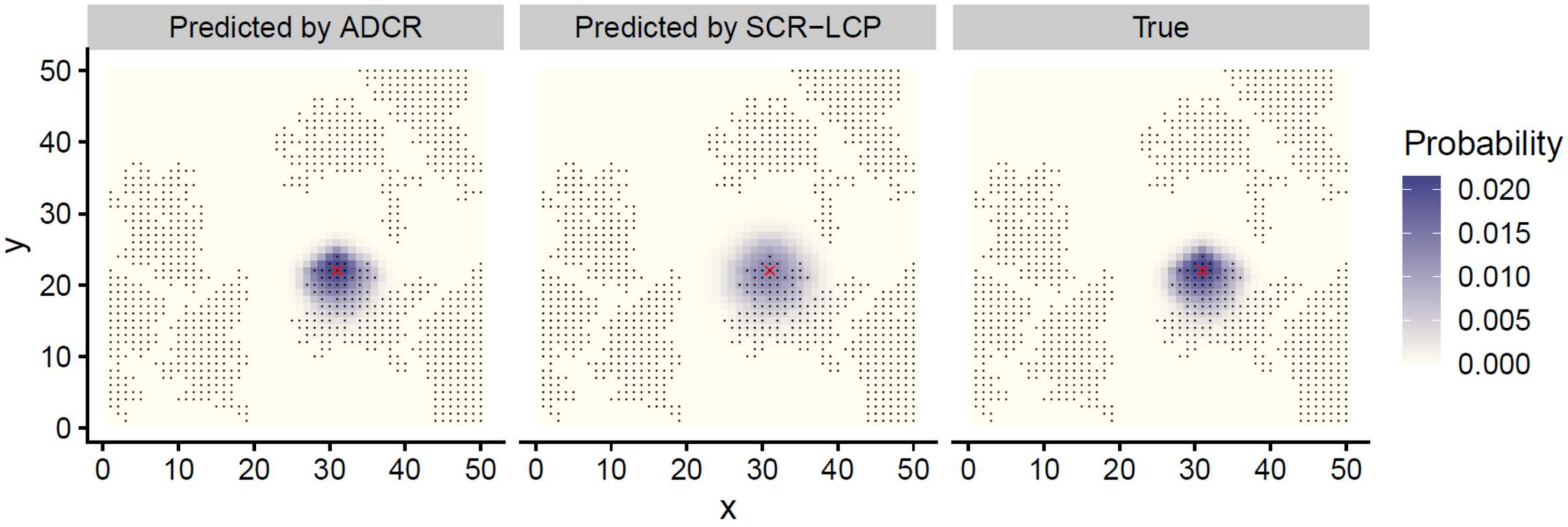

Fig. S5 Estimated log population density by ADCR, SCR with the least-cost path (SCR-LCP) and basic SCR under the scenario of severe data sparseness (i.e. $g_0 = -5$) and non-random detector alignment. Black dots are maximum likelihood estimates (MLEs). Open circles and and error bars indicate the mean and SD of MLEs. Horizontal black lines are the true value (-2.0).

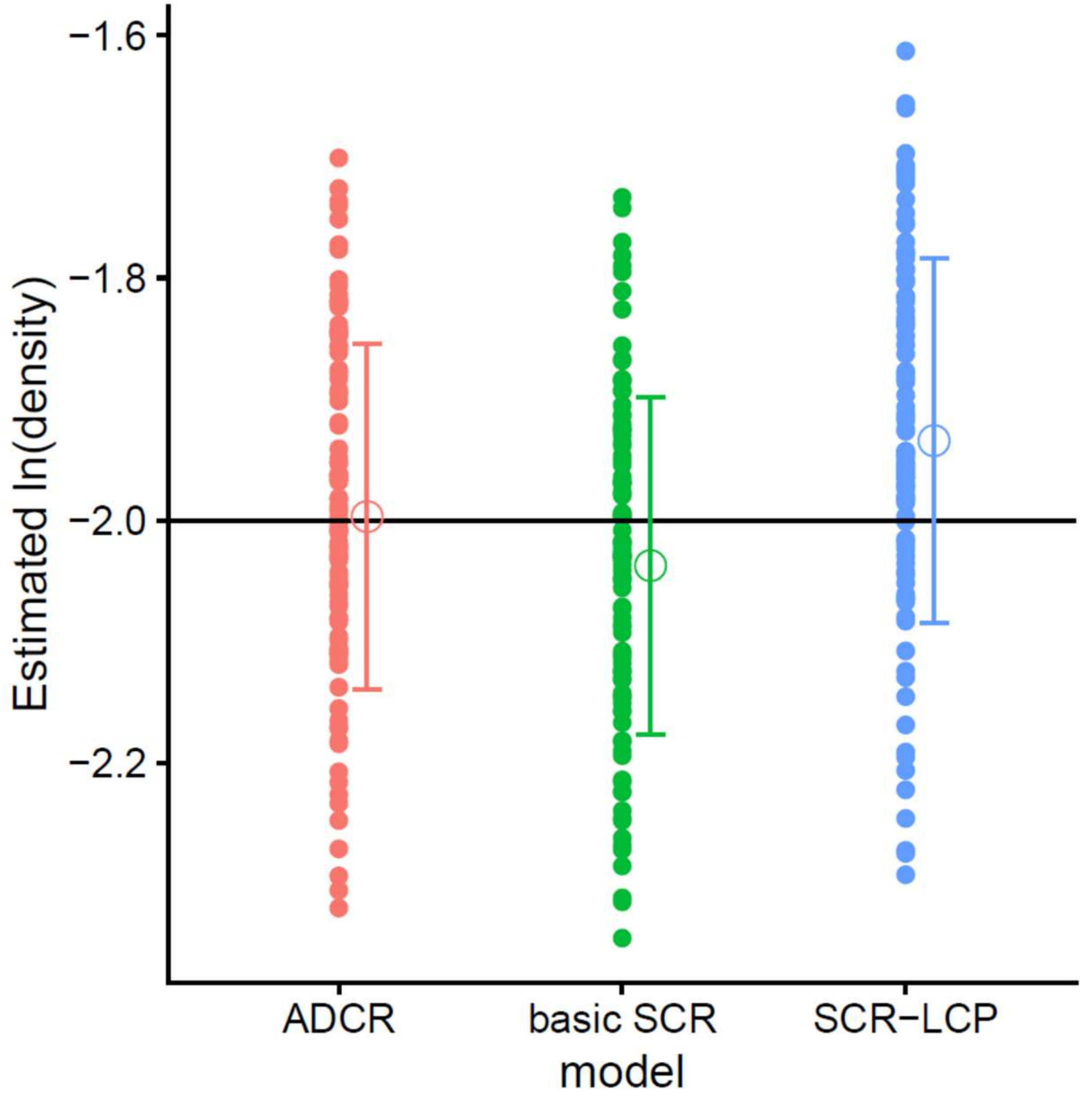

Fig. S6 Examples of predicted home ranges by (a) ADCR and (b) SCR with the least-cost path for the black bear. Red x symbols mark the home range centers. Black dots are cells contain water surface.

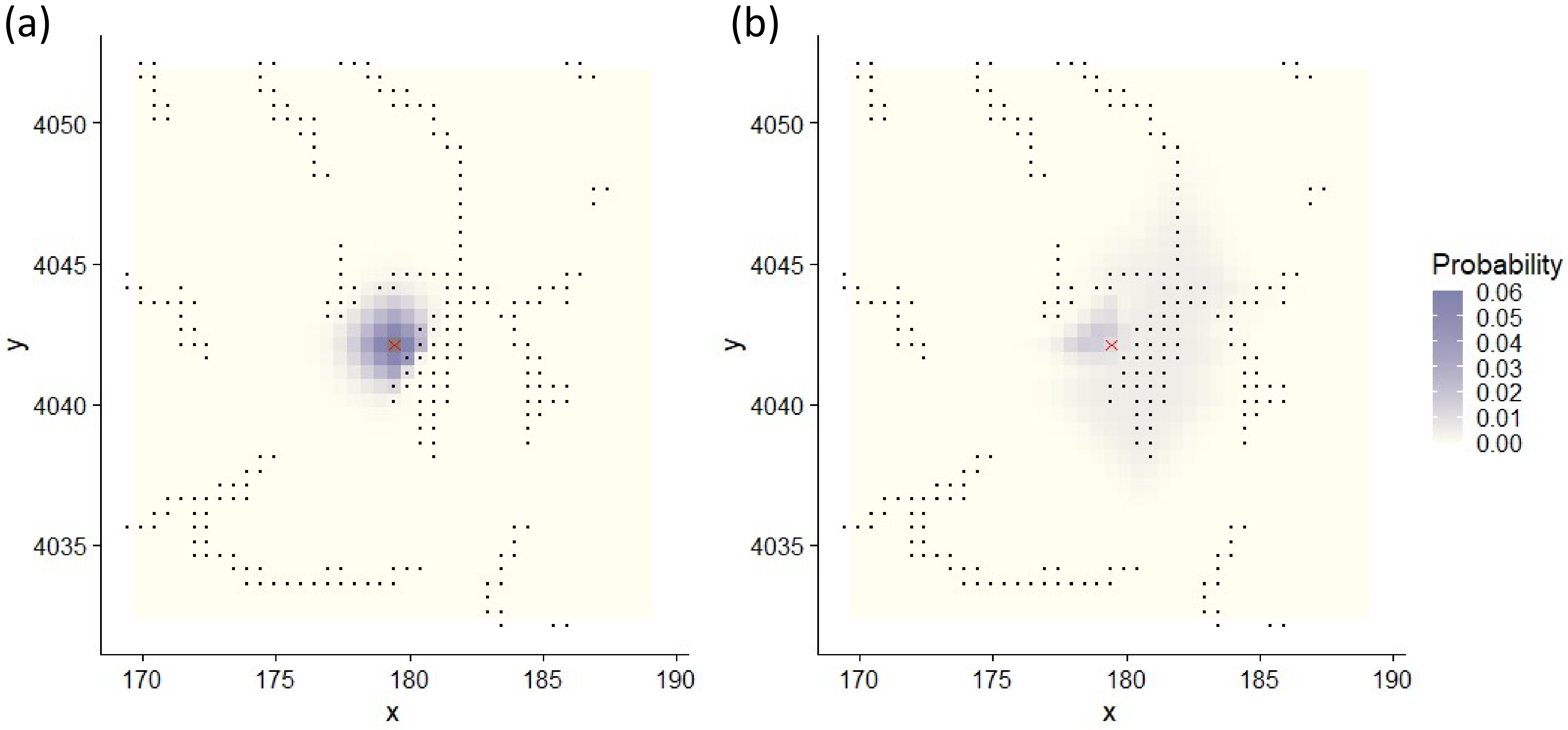

Fig. S7 Comparison between the estimates of baseline simulations and 1/2 resolution scenarios for (a) effects of landscape and (b) log population densities by ADCR. Dots and error bars are maximum likelihood estimates and 95% CI, respectively. Glay lines are principal components of parameter estimates.

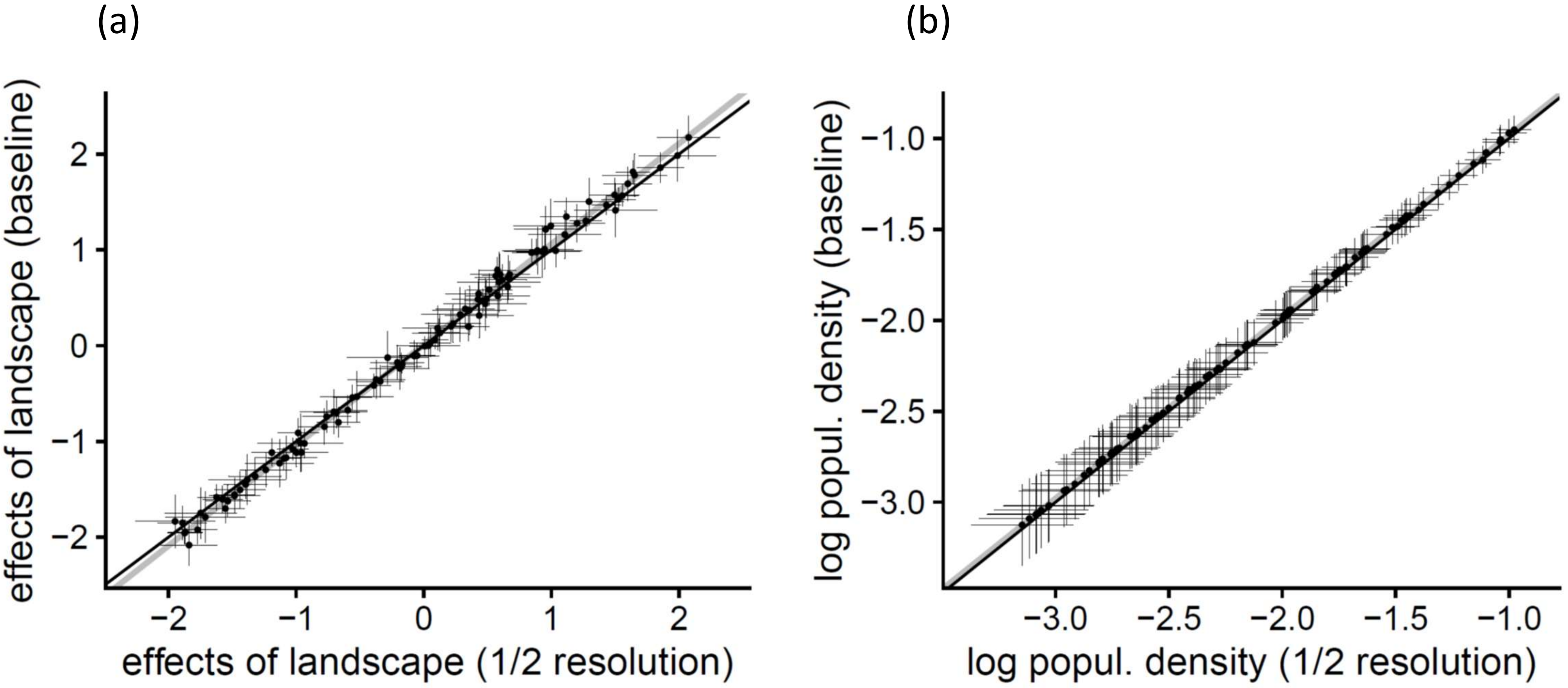
