## Appendix S6 for "Mechanistic home range capture–recapture models for the estimation of population density and landscape connectivity"

**Appendix S6** Supplementary Table.

**Table S1** Target parameter ranges for the neutral landscape generator, rflsgen. The parameter names were based on Justeau-Allaire et al. (2022). Parameters not shown were set to default values.

| Parameter name | Minimum | Maximum |
|---|---|---|
| Number of patches | 5 | 10 |
| Patch area | 100 | 400 |
| Total class area | 1000 | 2000 |
| Effective mesh size | 0 | 200 |

**Table S2** Contingency table on the correctness of sign (i.e. positive or negative) of landscape effect on connectivity estimated by ADCR and SCR with the least-cost path for the simulation scenario of sparse data and biased detector alignment. The significance was determined by non-overlap of 95% confidence interval with 0.

| | | effect on cost by SCR with the least-cost path | | | | |
|---|---|---|---|---|---|---|
| | | incorrect sign (significant) | incorrect sign (not significant) | correct sign (not significant) | correct sign (significant) | Total iterations |
| effect on permeability by ADCR | incorrect sign (significant) | 0 | 0 | 0 | 0 | 0 |
| | incorrect sign (not significant) | 0 | 1 | 1 | 0 | 2 |
| | correct sign (not significant) | 0 | 2 | 29 | 7 | 38 |
| | correct sign (significant) | 0 | 2 | 11 | 47 | 60 |
| | Total iterations | 0 | 5 | 41 | 54 | 100 |
